# First report of nematodes in *Lutzomyia edwardsi* (Diptera: Psychodidae, Phlebotominae) in cave, in the municipality of Sumidouro, State of Rio of Janeiro, Brazil

**DOI:** 10.1101/2020.09.20.305151

**Authors:** João Ricardo Carreira Alves, Cleber Nascimento do Carmo, Rodrigo Caldas Menezes, Mauricio Luís Vilela, Jacenir Reis dos Santos-Mallet

## Abstract

It is registered for the first time, the encounter of *Lutzomyia edwardsi* (Mangabeira) species of sandfly considered as a probable vector of *Leishmania* (*Viannia*) *braziliensis* (Vianna), main etiological agent of American cutaneous leishmaniasis in humans, for the municipality of Sumidouro, State of Rio of Janeiro. Leishmaniasis is currently considered a neglected tropical disease in the Americas. The objective of this study was to know the phlebotomine fauna in an area of Atlantic forest, with a cave in this municipality. From the sandflies collected, in 2009 and 2010, with a light trap, type CDC it is reported in a unique way, the encounter of a female *L. edwardsi*, infected with nematodes in a cave in the locality of São Caetano, in Sumidouro, state of Rio of Janeiro, Brazil. The description of sandfly and nematodes was carried out. The presence of this species in areas of leishmaniasis cases in the municipality of Sumidouro and its surroundings, requires the adoption of relevant entomological and / or epidemiological surveillance, by the Municipal Health Department of the region, when registering this disease.

---

A total of 1,008 species of sandflies are known worldwide, of which more than half are registered in the American continents. In Brazil, 279 species are known, representing 31% of the species known worldwide and with 22 proven or suspected species of leishmaniasis transmitting agents (Aguiar & Vieira 2003).

According to Young & Duncan (1994), in the New World, the subfamily Phlebotominae consists of the genera *Lutzomyia* França, *Brumptomyia* França and Parrot and *Warileyia* Hertig (Diptera: Psychodidae). The species of medical importance belong to the genus *Lutzomyia*.

From the discovery that some species of sandflies were vectors in the transmission of pathogens to humans and animals, studies on these insects have increased considerably, aiming to investigate their life cycle and identify which vectors occur in certain regions, considering the different species identified as important from an epidemiological point of view (Forattini 1973).

Carvalho *et al*. (2014) concluded from a literature review, based on collections made by the State Health Department of Rio de Janeiro, that the phlebotomine fauna of this state is composed of 65 species, eight of which belong to the genus *Brumptomyia* and 57 to the *Lutzomyia* genus. This research related the species, their distribution by municipality and reported that *Lutzomyia edwardsi* (Diptera: Psychodidae, Phlebotominae) was described by Mangabeira Filho in 1941 in the municipality of Nova Iguaçu, state of Rio de Janeiro. The specimens of this sandfly were collected in the forest, in a paca den (*Agouti paca L*.), in which seven males and two females were captured. In 1995, De Souza *et al*., Registered *L. edwardsi* in the municipality of São José do Vale do Rio Preto, later, Carvalho *et al*. (2014) pointed out, *L. edwardsi* in two municipalities in the mountain region: Bom Jardim and Petrópolis. Peres-Dias *et al*. (2016) researched the phlebotomine fauna of the municipality of Cantagalo, having collected 3,310 specimens from 12 species, eleven of the genus *Lutzomyia* and one of *Brumptomyia*. In this research, a male specimen of *L. edwardsi* was collected, being the first record of this species for this municipality.

In the mountainous regions of the state of Rio de Janeiro, from 2007 to 2017, 95 cases of American cutaneous leishmaniasis (ATL) were recorded, the highest occurrence being recorded in the municipality of Trajano de Moraes, with nineteen cases. Followed by Bom Jardim with nine cases. In Cantagalo, there were six cases from 2007 to 2015. In the municipalities of Duas Barras, Macuco, São Sebastião do Alto, Sumidouro and Teresópolis, one case in each unit (SINAN / SVS / WEB, 2019). In this region, four cases of visceral leishmaniasis (VL) were recorded, two in Petrópolis and two in Teresópolis (SINAN / SVS / WEB, 2019).

In order to report the encounter of *L. edwardsi* with nematodes, in addition to providing subsidies, which allowed the Health Departments of the above-mentioned municipalities to carry out a plan for the control and combat of leishmaniasis vectors, considering also that *L. edwardsi* has already been found infected with *Leishmania braziliensis* (Kinetoplastida: Trypanosomatidae) in the state of São Paulo. This work is justified and published.

The municipality of Sumidouro is located at Latitude 22° 2’43 “S, Longitude 42 ° 40’24” W, in the mountain region of the state of Rio de Janeiro, bordering the municipalities of Nova Friburgo, Teresópolis, Carmo, São José do Vale do Rio Preto, Sapucaia and Duas Barras (Fig 1). The study area was described in more detail by Alves *et al*. (2020)

**Fig. 1.**
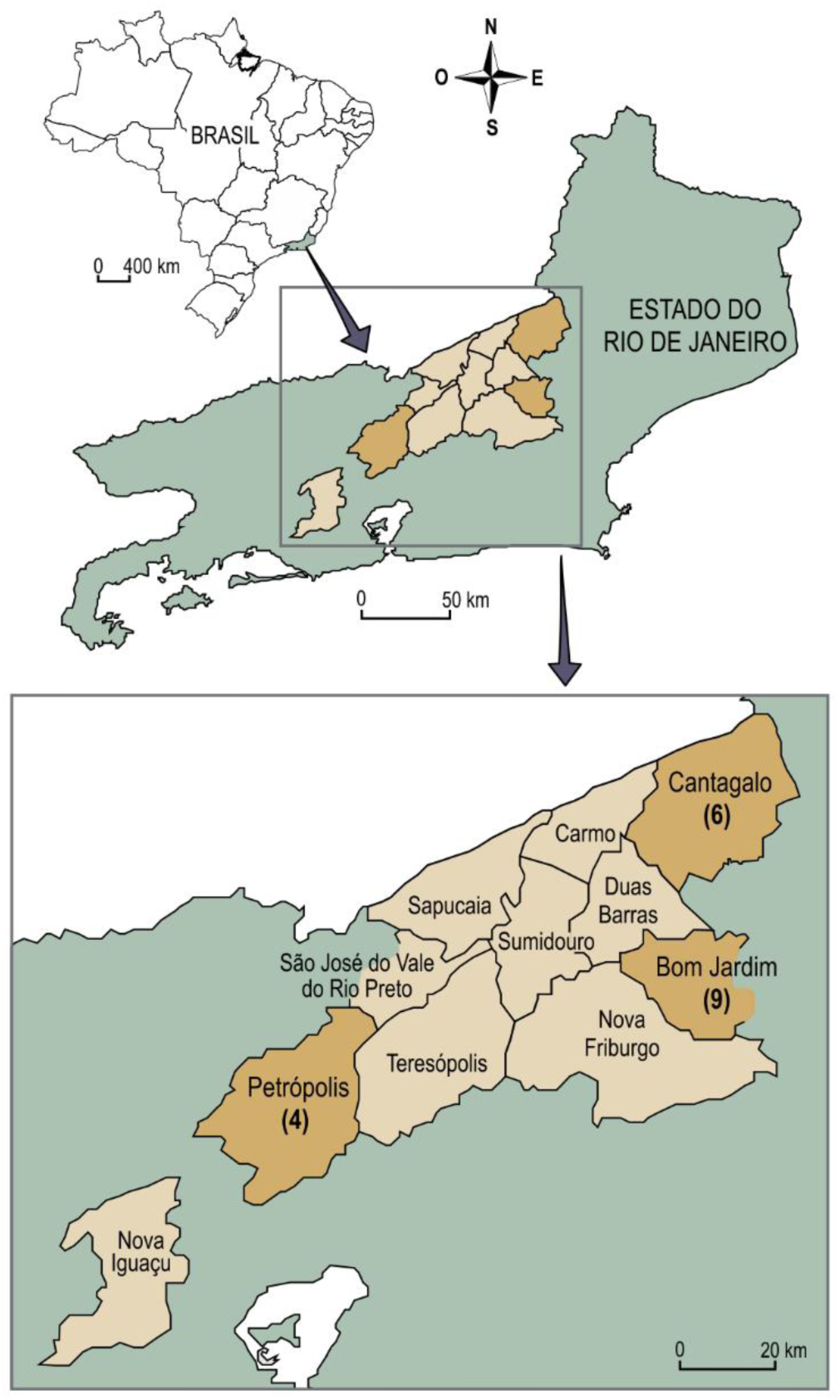
Geographic location of the neighboring municipalities of Sumidouro: Carmo (21° 55’ 53. 40’’ S / 42° 36’ 18. 23’’ W), 2 Duas Barras (22° 03’ 15. 60’’ S / 42° 31’ 24. 24’’ W), 3 Nova Friburgo (22° 17’ 19. 58’’ S / 42° 32’ 02. 88’’ W), 4 -Teresópolis (22° 25’ 05. 35’’ S / 42° 58’ 15. 49’’ W), 5 São José do Vale do Rio Preto (22° 13’ 00’’ S / 42° 51’ 06’’ W), 6 Sapucaia (21° 59’ 40’’ S / 42° 54’ 57’’W), 7 Sumidouro (22° 2’43 “S / 42 ° 40’24” W). 8-Nova Iguaçu (22° 45’37.16” S / 43° 26’51.82” W) Locality type of *L. edwardsi*. The number in parentheses corresponds to the cases of leishmaniasis registered in the municipalities of Bom Jardim, Cantagalo and Petrópolis from 2007 to 2017, where *L. edwardsi* was found.

Poisson regression was performed to assess and prove that the collections of *L. edwardsi*, between the São Caetano cave, its surroundings and the forest, showed statistically significant differences.

In order to describe the phlebotomine and the entomoparasite, the Zess microscope, with Axiolmager, equipped with Apo Tome, was used to perform the photos, measurements and formatting of the nematodes and the specimen of *L. edwardsi*. (Fig 2 a, b, c, d)

**Fig 2.**
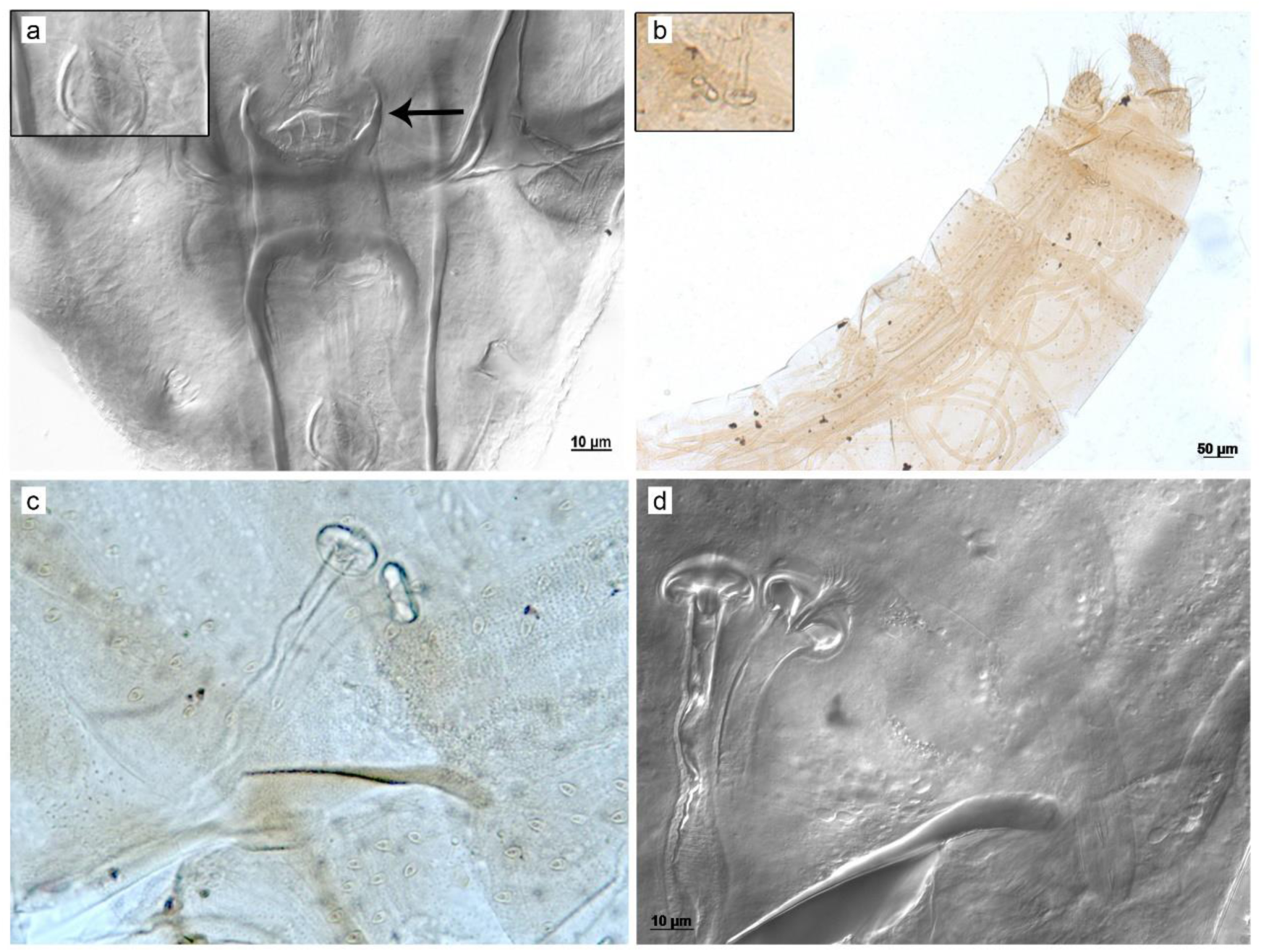
a-Cibary of the female of *L. edwardsi*, with vertical teeth, in detail shows a nematode found on the head. b -Abdomen with nematodes, *L. edwardsi* spermateca. c -Details of the spermatec in bright field, d -Individual duct, body and head.

The identification was made using the taxonomic key of Poinar *et al*. (2002), and through the photos of the nematodes that were sent to Dr. Poinar Jr, Department of Entomology; Oregon State University, United State of American.

1,756 sandflies belonging to 14 species were captured, 11 of which from the genus *Lutzomyia*: *Lutzomyia gasparviannai* Martins, Godoy & Silva, *L. edwardsi* (Mangabeira), *L. tupynambai* (Mangabeira), were the most prevalent and three species of the genus *Brumptomyia*. The methodology, the material used and the results are described by Alves *et al*. (2020).

A total of 290 specimens of *L. edwardsi* were collected, 67 males and 223 females. In the cave, there were 37 males and 114 females, among these specimens, it was found the existence of a female of *L. edwardsi*, infected with more than 11 nematodes in the abdomen.

Poinar et al. (1993) found in a colony of *L. longipalpis*, in Colombia, specimens infected with nematodes, which were taken to a laboratory in the United States. The authors described *Anandranema phlebotophaga* n. gen., n. sp. (Allantonematidae: Tylenchida).

The nematodes described by the authors are morphologically similar to the nematodes found in our study. Poinar *et al*. (1993) also point out that these nematodes are making the females of *L. longipalpis* infertile, leading the authors to consider them an entomoparasite and suitable to be used as biological control for this species.

In 2001, in the municipality of Cotia, state of São Paulo, Brazil, a case of visceral leishmaniasis occurred in a cat. During the research, infection of *L. edwardsi* by *Le. (Viannia) braziliensis* was recorded, by PCR, etiological agent of American cutaneous leishmaniasis.

Which led us to consider its relevance in the studied area, since two cases of ATL were registered in the region. In addition, the species was collected at the three collection sites, demonstrating a wide distribution in the studied location. It is noteworthy that the region is frequented by farmers, residents, hunters, students, paleontologists, biologists.

In 2002, Poinar *et al*. describe a new nematode, *Elaeolenchus parthenonema* n. gen., n. sp. (Nematoda: Sphaerularioidea: Anandranematidae n. Farm.) In an *Elaeidobius kamerunicus* (Curculionidae: Coleóptera). The new genus is placed in Anandranematidae n. farm., together with the genus *Anandranema* Poinar *et al*. (1993), as they are characterized by being nematodes with a single auto generation.

The authors confirm the genus and species of *Anadranema phlebotophaga*, which resembles the nematodes found by us.

In 2002, Secundino *et al*. detected and morphologically characterized a new entomoparasite of *L. longipalpis*, using electron microscope, fluorescence and differential. They point out that the adult forms of these nematodes are different from those described by Poinar *et al*. (1993) and those found in Sumidouro. Studies are being done to identify and describe it.

*L. edwardsi* were collected in São José do Vale do Rio Preto, Bom Jardim, Cantagalo and Petrópolis, the last three municipalities were responsible for 17 cases of ATL, between 2007 and 2016. In Petrópolis, two cases of VL were registered, in 2016.

In absolute terms, the highest number of *L. edwardsi* occurred in Sumidouro, considering the cases of ATL in these four municipalities, the number of females was more accentuated in our study. Catches with human bait were made in Petrópolis, with negative results for this species. The methodology used in Bom Jardim differs from that applied in the other three municipalities, in which light is used to attract sand flies. It is evident that the CDC trap, with light bait, is more efficient, whether in the home, in the woods or in the forest. However, inside and outside a house and / or shelter for domestic animals in an area with human cases, the castro catcher is recommended. These works, show us that *L. edwardsi* can be found in a cave environment, in its surroundings, in the forest, and in the state of Rio de Janeiro.

The results of the Poisson regression showed that there was a significant difference (p <0.01) between the collections of the cave and the forest, as well as, of the forest and the surroundings for *L. edwardsi*. Which confirms the importance of the collections made in Sumidouro.

Galati *et al*. (2010) found *L. monticola* infected with microfilarias in the Speleological Province of Vale do Ribeira, São Paulo. In his article, he concludes that this species is anthropophilic.

In this study, *L. monticola* was not found, which does not corroborate Galati *et al*. (2010), nor microfilarias and there is no evidence that *L. edwardsi* is anthropophilic, which differs from Souza *et al*. (2002). Which suggests that the nematodes found could not be transmitted to man.

In 2011, Souza *et al*. recorded the encounter of *L. fischeri* infected by nematodes belonging to the family Allantonematidae and the genus *Anandranema*. This material was found in the municipality of Rio de Janeiro, state of Rio de Janeiro.

In this research *L. fischeri* was not found, however, Carreira-Alves (2008) registered this species in the municipality of Carmo. According to the literature, this species is anthropophilic, wild and is adapting to the anthropic environment. (Aguiar & Vieira, 2018) Preliminary studies on nematodes highlight the difference between the size of the stylus, the position of the gonads in relation to that described by Poinar *et al*. (1993), as well as the existence of the cuticle, in the anterior part, in young parasites (Fig 3 a, b, c) (Fig 4 d). It is also noted that some species are also hermaphrodites in agreement with the others cited by the author (Fig 4 d). These observations were made according to the taxonomic key of Poinar *et al*. (2002).

**Fig 3.**
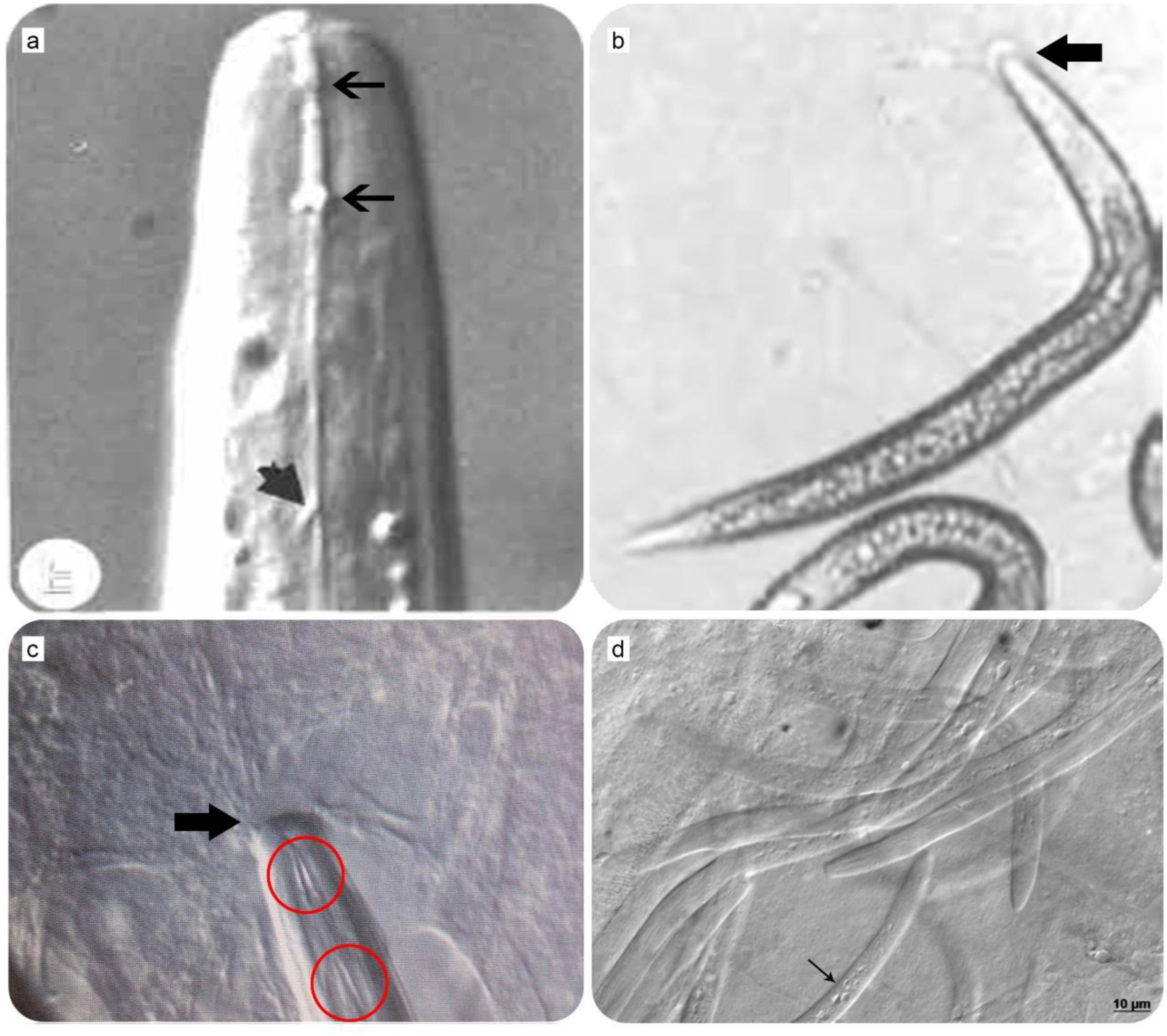
a and b -*Anandranema phlebotophaga* species described by Poinar *et al*, 1993. c and d - are the species that were collected in this work. The red circle and a thick black arrow mark the stylets (a and c), while the arrows (b and c) show the cuticles. Thin black arrows -indicate the sub -ventral opening of the gonads (a). Fig. 4 -According to Poinar, some of these species are hermaphrodites. (d).

The catch of *L. edwardsi*, infected with nematodes, is registered for Brazil. Considering that they are hermaphrodites, have stilettos, anterior gonads, and are entomoparasites of sandflies, as the literature points out. We believe that they belong to the Family Anandranematidae n. fam.

Because *L. edwardsi* and the nematodes were mounted between slide and cover slip with berleze, it was not possible to identify even the species of the nematodes

The fact that it was collected in a cave in areas with stones and forest of volcanic origin, differentiates it from the reports of the aforementioned authors.

The existence of this species in the area of cutaneous leishmaniasis cases, must have a more conscious and effective entomological and / or epidemiological surveillance, by the Municipal Health Department of the region. Thus, allowing this information to be used for the study and development of technical standards for the control and prevention of vectors and leishmaniasis.

## Acknowledgements

To Dr. Felipe Ferraz Figueiredo Moreira and Dr. Márcio Felix, Laboratory of Entomological Biodiversity of Fundação Oswaldo Cruz for logistical support. Dr. Francisco Gerson de Araújo, Federal Rural University of Rio de Janeiro for support and learning. The Heloisa Maria Nogueira Diniz the production and processing of images. Coordination for the Improvement of Higher Education Personnel -Brazil (CAPES) -Funding Code 001.

